# Spatial organization of collective food distribution in a paper wasp society

**DOI:** 10.1101/2023.10.13.562279

**Authors:** Nitika Sharma, Raghavendra Gadagkar

## Abstract

In social insect colonies, food transferred through space and time via nestmates carries both nutrition and information. We study the mechanism of spatio-temporal coordination (or the lack thereof) between multiple wasps for them to optimally solve the problem of feeding randomly placed larvae in a social insect colony. We followed each morsel of food brought into semi-natural colonies of the tropical paper wasp *Ropalidia marginata,* in each of 36 feeding bouts until the food was exhausted. We found that most acts of feeding larvae are performed by wasps that unload food from foragers, but unloading itself is highly skewed among individuals. We also found that larvae closer to the nest center were fed more frequently than those at the periphery, irrespective of their developmental stage. This differential feeding may have fitness consequences as it is known that well-fed larvae become voracious feeders as adults and have a higher likelihood to become reproductives. Using the analogies of the travelling salesman problem and the Hamiltonian path problem, we showed that individuals which disburse food to more larvae within a feeding bout adopted paths that were considerably shorter than expected by chance while the distinction between adopted and random routes was not as pronounced when the number of larvae fed in a bout was fewer. There was no spatial segregation between wasps feeding larvae in parallel, possibly building redundancy and avoiding larval starvation across bouts. Understanding the spatial organization of food transfer may be a key to understanding how insect societies achieve efficient social organization and division of labor.

**Significance:** The limited studies on within-nest spatial organization of food distribution have been performed on ants and not primitively eusocial insects like wasps that contain randomly placed yet trackable larvae within the colony. We address the gap in a holistic understanding of spatial organization of food distribution through our study in the social wasp *Ropalidia marginata*. Through the lens of task partitioning, colony-level larval landscape, and spatial strategies adopted at individual and collective scales, our study presents one of the most detailed account of behavioral and spatio-temporal data mapping sources and sinks of food in a colony chronologically. We found that feeding routes adopted by wasps while feeding randomly placed larvae within a colony may not be a result of random walk but instead the degrees of route optimization based on the food load available to distribute within each feeding bout. We suggest paper wasp colonies to be ideal model systems for spatial studies owing to their observable colony sizes and randomly placed brood. Understanding the mechanism of colony-level problem solving by individuals with limited local information has widespread real-life applications besides a better understanding of social organization in colonies of social insects.

## Introduction

Social insects have had unmatched ecological success, possibly due to their collective abilities to solve problems efficiently as superorganisms (Wilson, 1990). While no single individual in the colony has the cognitive capacity of the collective, the colony as a whole is more than the sum of its parts and regularly solves ecological problems and makes optimal decisions using simple rules (Camazine, 1991). Thus, colony functioning is decentralized such that no single individual has global knowledge about the colony’s needs. Instead, the colony is comprised of individuals with limited local information who cooperate to fulfil those needs by following simple, local rules. Individuals in social insect colonies are often redundant, and the colony continues to function normally even when some of them perform incorrectly or die (Hirsh and Gordon 2001). Colony needs may range from deciding which nest to migrate to, allocating tasks among colony members, food acquisition and distribution. Emergent colony-level behaviors based on the distributed organization of individuals have been well-studied in social insect colonies in the contexts of foraging (Deneubourg et al. 1986; Nicolis and Deneubourg 1999; Dorigo et al. 2000), nest building and searching (Theraulaz et al. 1999; Pratt et al. 2002; Pratt 2005) and task allocation (Gordon 1992, 1996; Pacala et al. 1996). However, the mechanisms of food distribution among colony members and brood remain poorly studied.

Food processing and distribution are among the most conspicuous and coalescing tasks of a social insect colony that bind all colony members in a “food circulatory system” irrespective of age, caste, and reproductive roles (Meurville & LeBoeuf, 2021). Food exchange in some social insects is also accompanied by the exchange of information about food sources, food odor (Sola & Josens, 2016), gut microbiota (Lanan et al., 2016), defence-related molecules to fight infection (Hamilton et al., 2011), hormones that determine larval developmental fate (LeBoeuf et al., 2016; Walsh et al., 2018), etc. Typically, an arriving forager unloads food to many intra-nidal (within colony) workers, who then distribute it among other adults and the larvae. This multi-level food distribution also dilutes any toxic substances brought in by the foragers (Greenwald et al., 2019). The higher rates of intra-caste interactions compared to inter-caste interactions also limit the rapid spread of pathogens in social insect colonies (Mersch et al., 2013). Such differential social interactions are often the result of non-random spatial distribution of individuals within social insect colonies (Heyman et al., 2017; Mersch et al., 2013; Sharma & Gadagkar, 2019). Younger individuals that tend to nurse the brood in colonies are generally found closer to the colony center, while older individuals that function as foragers are located towards the periphery (Jandt & Dornhaus, 2009; A.B. Sendova-Franks & Franks, 1995). The spatial location of a worker automatically decides who she interacts with (Jeanson, 2012).

Despite the wide-ranging contexts and implications of food exchange within colonies, few studies have examined it as a distributed problem solved collectively by many individuals. While network-based studies of trophallactic interactions can help quantify colony performance in terms of robustness and resilience (Bles et al., 2018; Quque & Bles, 2021), they often overlook the mechanism of food distribution embedded in colony space (but see (Aurélie Buffin et al., 2009; Ana B. Sendova-Franks et al., 2010)). The mechanism of spatial food distribution in social insect colonies has only recently begun to receive attention after the development of advanced tracking methods (Greenwald et al., 2015). The handful of existing studies on the spatial aspect of food distribution (Greenwald et al., 2019; Planckaert et al., 2019) exclusively record adult-adult food exchange spatial locations and ignore the spatial locations of adult-larval interactions because the latter are harder to observe (Meurville & LeBoeuf, 2021). But larvae are a critical component of the food network as sinks of the food brought into the colony, and their location within the colony may dictate decisions about food distribution. Furthermore, no study in our knowledge has attempted to understand the mechanism of food distribution by multiple individuals working simultaneously.

The few studies that spatially track each trophallactic event and the role of each individual in food dissemination are confined to ant or honey bee colonies that have clustered brood whose individual locations are variable (Camazine 1991; Feinerman and Korman 2013; Greenwald et al. 2015, 2018; Planckaert et al. 2019). Nests of bumble bees and wasps offer opportunities to individually track the feeding frequency of larvae located in distinct cells (Strassmann et al. 2000; Couvillon and Dornhaus 2009). The nests of paper wasps are an ideal system to study the exchange of food among adults and between adults and larvae because of their relatively small colony sizes and the presence of larvae in discrete cells.

In the present study, we focus on the distribution of food on the nests of the tropical primitively eusocial paper wasp *Ropalidia marginata*. The brood on the nests of this species is located in discrete cells randomly located on the nest surface, irrespective of their developmental stage (Sharma and Gadagkar 2019). Colonies consist of a single egg-layer (queen) and several non-egg-laying workers. Brood is present on the front of unenveloped nests, making it easy to observe feeding of larvae by the adults and trophallactic interactions between adults and larvae (Gadagkar 2009). The foragers in our observation colonies were free to fly out and bring back prey from their natural environment and are thus expected to be closer to natural colonies in terms of frequency of forager arrivals and the amount of food brought in. Some previous studies have additionally shown that the frequency of feeding the larvae is a better proxy for the quantity of food exchanged than the time duration of each trophallactic event (A. Buffin et al., 2011). The distribution of food by adults to the randomly located larvae is a complex problem; in this paper, we attempt to understand how these primitively eusocial wasps collectively solve it. In particular, we study how the task of food distribution is organized among colony members. We ask whether individuals preferentially feed certain larvae over others and how multiple individuals distributing food in parallel within a feeding bout spatially organize their feeding effort and navigate to reach the randomly located larvae.

## Results

Here we present our analysis of 36 complete feeding bouts observed on three nests of the tropical primitively eusocial paper wasp *Ropalidia marginata*. We consider as one feeding bout, all events of food exchange among adults and between adults and larvae, starting with the arrival of a forager on the nest up to the exhaustion of the food she has brought. If a second forager arrives with food before the food brought by the first forager is exhausted, we extend the duration of the bout until all the food is exhausted. In a feeding bout, the number of foragers ranged from 1 to 3, the number of primary receivers ranged from 1 to 10 and the number of secondary or tertiary receivers from 1 to 9. Feeding bouts lasted 6.16 ± 4.85 minutes (mean ± sd; range = 0.78 – 19.28). Within a feeding bout, 3.94 ± 2.63 wasps (range 1 to 13) simultaneously distributed food to the larvae and 22.8% ± 15.3% (range: 1.9% - 70.2%) of the larvae on the nest were fed. As expected, the number of feeders distributing food to the larvae was strongly correlated with the total number of larvae fed in a bout (Kendall’s correlation coefficient = 0.66, p < 0.01). In 13 out of 36 bouts, more than one forager arrived on the nest with food at intervals of 2.54 ± 1.89 minutes (Fig. S4).

*Task Partitioning.* Some wasps (primary receivers) unload food from incoming foragers and distribute it to other wasps (secondary or tertiary receivers). Foragers, and the primary, secondary and tertiary receivers all fed larvae to different extents (Fig. 1). In a bout, only 16%± 10.4% (range = 4% to 50%) of the wasps that brought or received food fed larvae while the remaining consumed the food, transferred it to other adult wasps or fed larvae in future bouts. Within a feeding bout, 23.9% ± 8.9% (range = 3.8% to 41.0%) of the wasps acted as primary receivers, 16.4 %± 10.1% (range = 0% to 41.0%) acted as secondary or tertiary receivers. In the three nests, some foragers (58.8%, 45.5%, 25.0%) fed larvae with or without unloading some of the food to other adult wasps, while others (41.2%, 54.5%, 75.0%) did not feed larvae and only unloaded food to the adult wasps. We measured the extent of task partitioning between bringing food and feeding larvae by computing the Gorelick’s division of labor index. We obtained values of 0.39, 0.62 and 0.39 for the three nests respectively. These values were significantly greater than expected if wasps randomly performed either task (see Fig. S5).

**Fig. 1:**
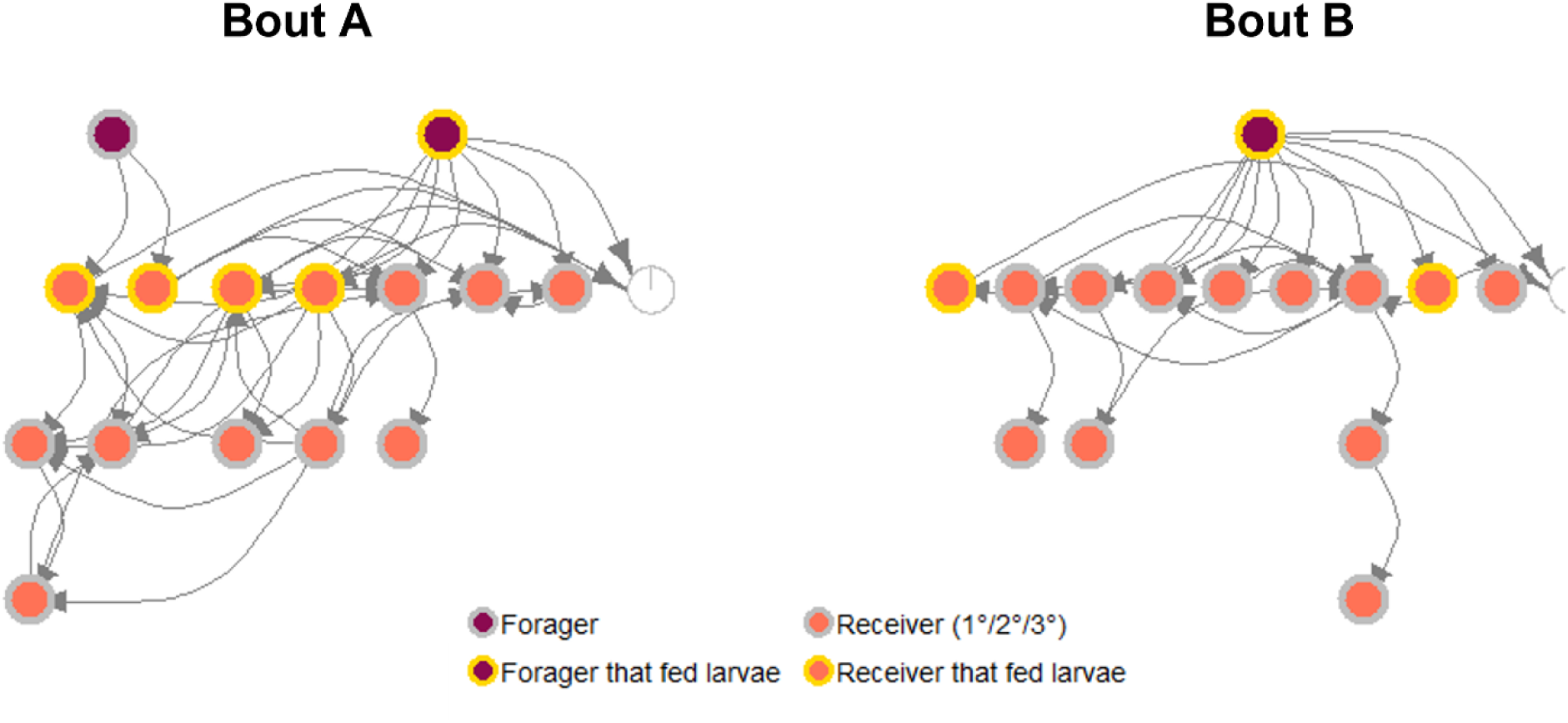
Schematic representation of a simple and a more complex feeding bout. Note that in bout A for example larvae were fed by both the forager and by two primary receivers while six primary receivers did not feed larvae, while two of them transferred some food to secondary receivers who also did not feed larvae.

Queens were intermediate (as compared to workers) in terms of receiving food (either from foragers or from the receivers). When queens received food (15 instances in 36 bouts), they usually consumed it themselves and only very rarely fed larvae (only once in 36 bouts) or lost the food to other wasps (in 14 out of 36 bouts) (Fig. S6). Of the primary receivers, 38.9% ± 23.4% did not feed larvae, 41.6% ± 31.4% fed larvae, while 19.8% ± 20.7% acted as ‘conduits’ i.e., transferred the food to other adult wasps. The task of unloading food from the foragers was not performed equally by different workers. We measured inequality in the performance of this task by computing the Gini coefficient which is normally used by economists to measure wealth inequality such that 0 indicates complete equality and 1 indicates complete inequality. We obtained values of 0.49, 0.44 and 0.52 respectively for the three nests, suggesting considerable departure from equal participation (Fig. S8).

### Differential feeding of larvae

Of 142 instances of a wasp receiving food and feeding larvae in a bout, in 33 cases feeders were observed to inspect but not feed some larvae (2 ± 2.2 larvae) previously fed by another feeder in the same bout. 31% of the colony members displayed such skipping of previously fed larvae. In 57 of the 142 cases, some larvae (2 ± 1.2 larvae) were skipped even though they were not previously fed in that bout. These unfed larvae were later fed by another feeder in the same bout. The number of times a larva was fed was significantly influenced by its stage of development such that larger larvae were fed more often (linear mixed effects model with Poisson error family, Fig. S9 and Table S2). Moreover, irrespective of the stage of development of larvae, the cells closer to the colony centre were fed more often than those at the periphery of the nest (Fig. 2, S10 and Table 1).

**Fig. 2:**
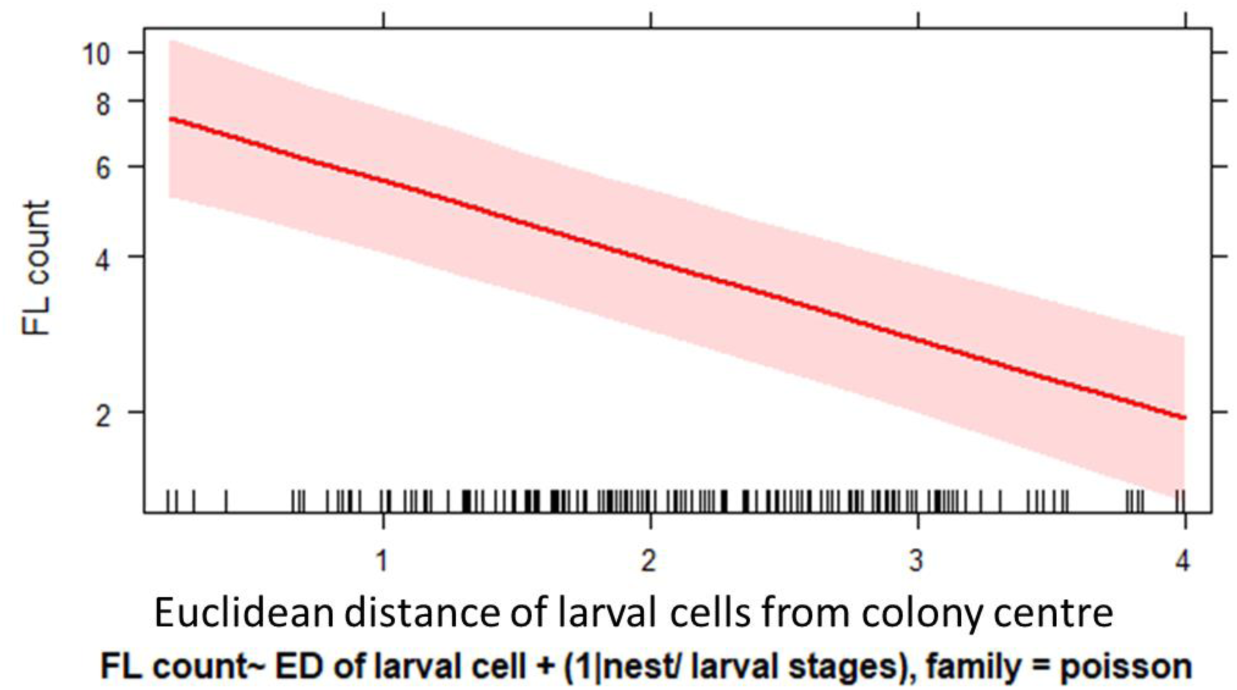
The larval cells closer to the colony centre were fed more number of times than the ones at the periphery. Linear mixed effects model with larval stages nested in nest ID as random effect and Poisson error family.

**Table 1:** The larval cells closer to the colony centre were fed more number of times than the ones at the periphery. Linear mixed effects model with larval stages nested in nest ID as random effect and Poisson error family.

|  | Estimate | Std. error | z value | CI (2.5%) | CI (97.5%) | Pr(>\|z\|) |
| --- | --- | --- | --- | --- | --- | --- |
| Intercept | 2.07 | 0.18 | 11.46 |  |  |  |
| ED of larval cell | -0.35 | 0.04 | -7.9 | -0.44 | -0.26 | **<0.001**<br>*** |

### Spatial organization of larval feeding - all bouts pooled

To evaluate space use during food distribution, we delineated for each feeder (considering all her feeding bouts), the area on the nest where she concentrated 50% of her feeding efforts. We label this as the core feeding area. Such feeding core areas occupied 16.78% ± 7.90% of the nest (see Fig. 3 for example). The overlap of the core feeding areas of the different wasps was less than expected by chance (Figure 3B).

**Fig. 3:**
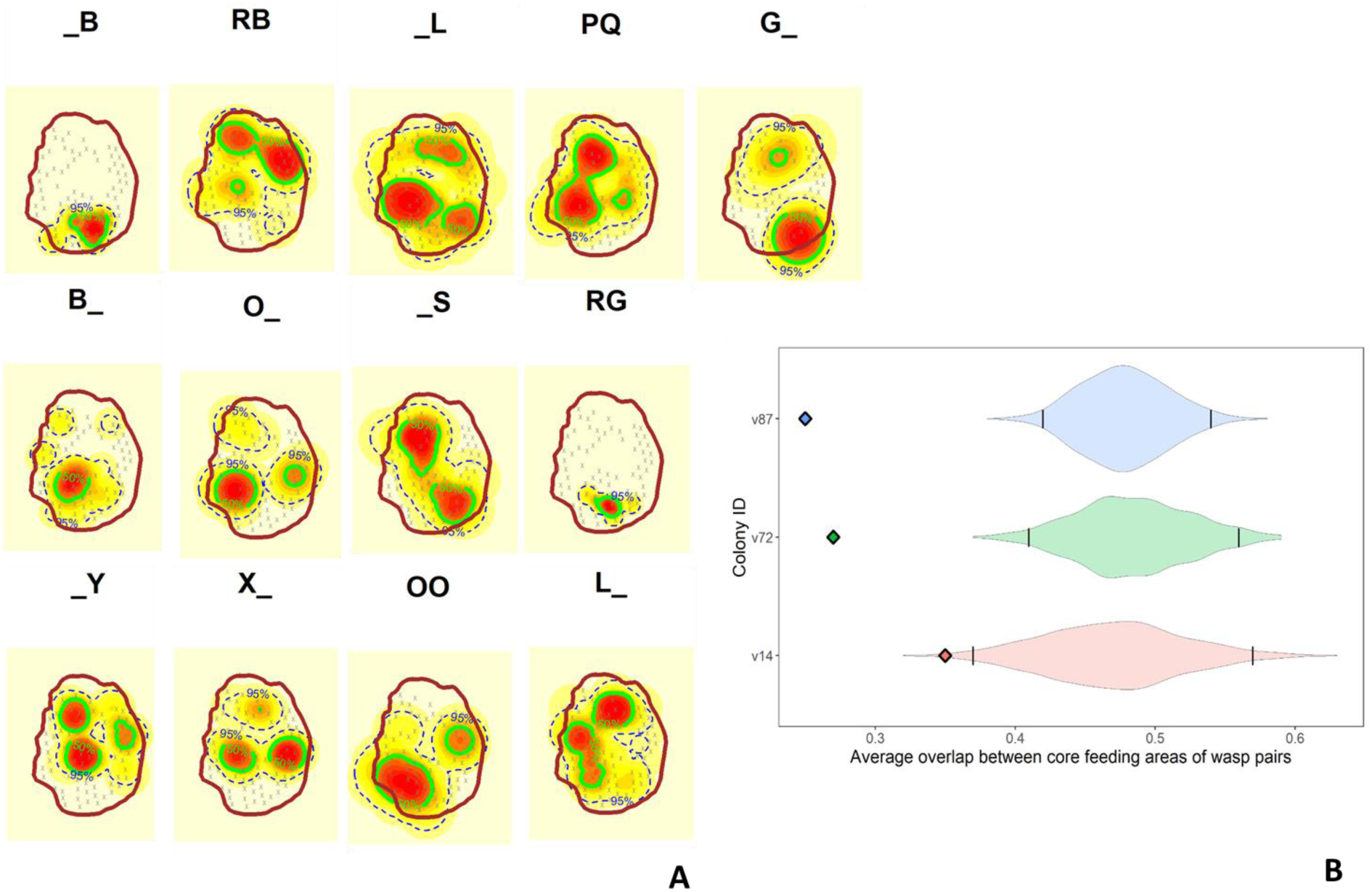
A) 50% core feeding areas of all feeders: the area of frequent feeding (50% core area = green solid, 95% range = blue dashed, larval locations = gray crosses) by each of the feeders over three days on the nest v87. B) The average overlap of feeding core areas between pair of wasps observed on a nest (points in each row) was significantly lesser than expected from random larval feeding (simulated distribution as violin plots). The confidence interval of simulated overlaps is denoted as vertical black lines on the violin plots.

### Spatial organization of larval feeding - within bouts

The median proportions of larvae fed by a single wasp in a bout were 0.89, 0.89 and 0.71 for the three nests. These numbers were not significantly different from those expected by chance as revealed by simulations. The corresponding median proportions in simulations where wasps fed larvae randomly were 0.87, 0.89 and 0.81 (Wilcoxon test, W=48616, p = 0.4; W = 85310, p = 0.5; W = 33491, p = 0.9). We therefore conclude that in a feeding bout, different feeders did not avoid feeding the same larvae (Fig. 4).

**Fig. 4:**
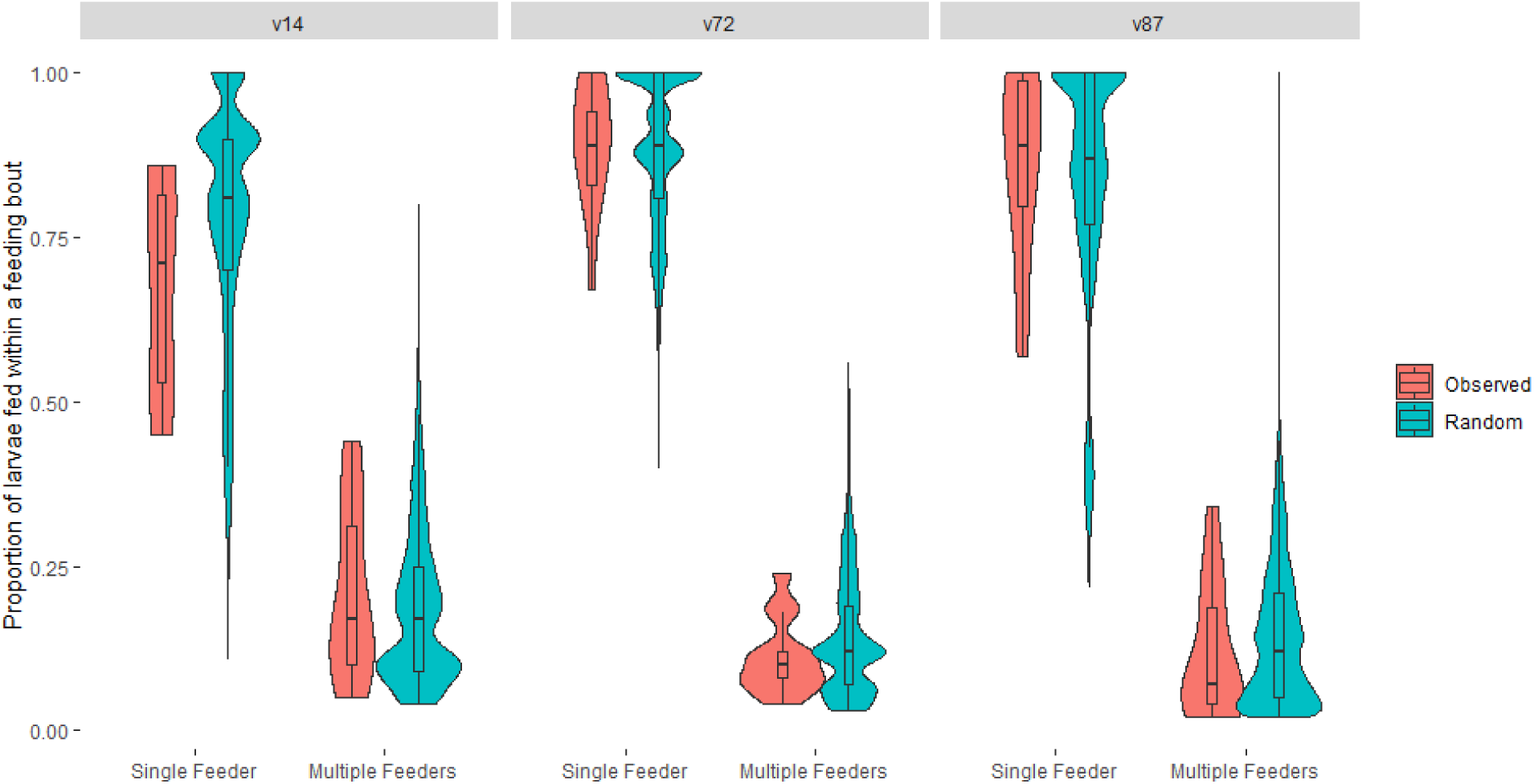
The observed proportion of cells fed by a unique wasp within feeding bouts was not different from simulations assuming no spatial avoidance between simultaneously feeding wasps in a feeding bout.

### Spatial organization of larval feeding - feeding routes

The number of larval cells fed by an individual feeder in a feeding bout ranged from 1 to 23 cells (7.32 ± 4.82). We have spatial data on 81 complete feeding routes of wasps that fed at least three larvae. Of these, 27 routes (33.3%) were shorter than expected by chance (see Fig. 5 and S11 for examples). The ratio of the normalized length of the expected route to the normalized length of the observed route increased significantly with the number of larvae being fed. Thus, feeders used routes that were shorter than expected especially when they had to feed many larvae in a bout (Fig. 6 and Table 2).

**Fig. 5:**
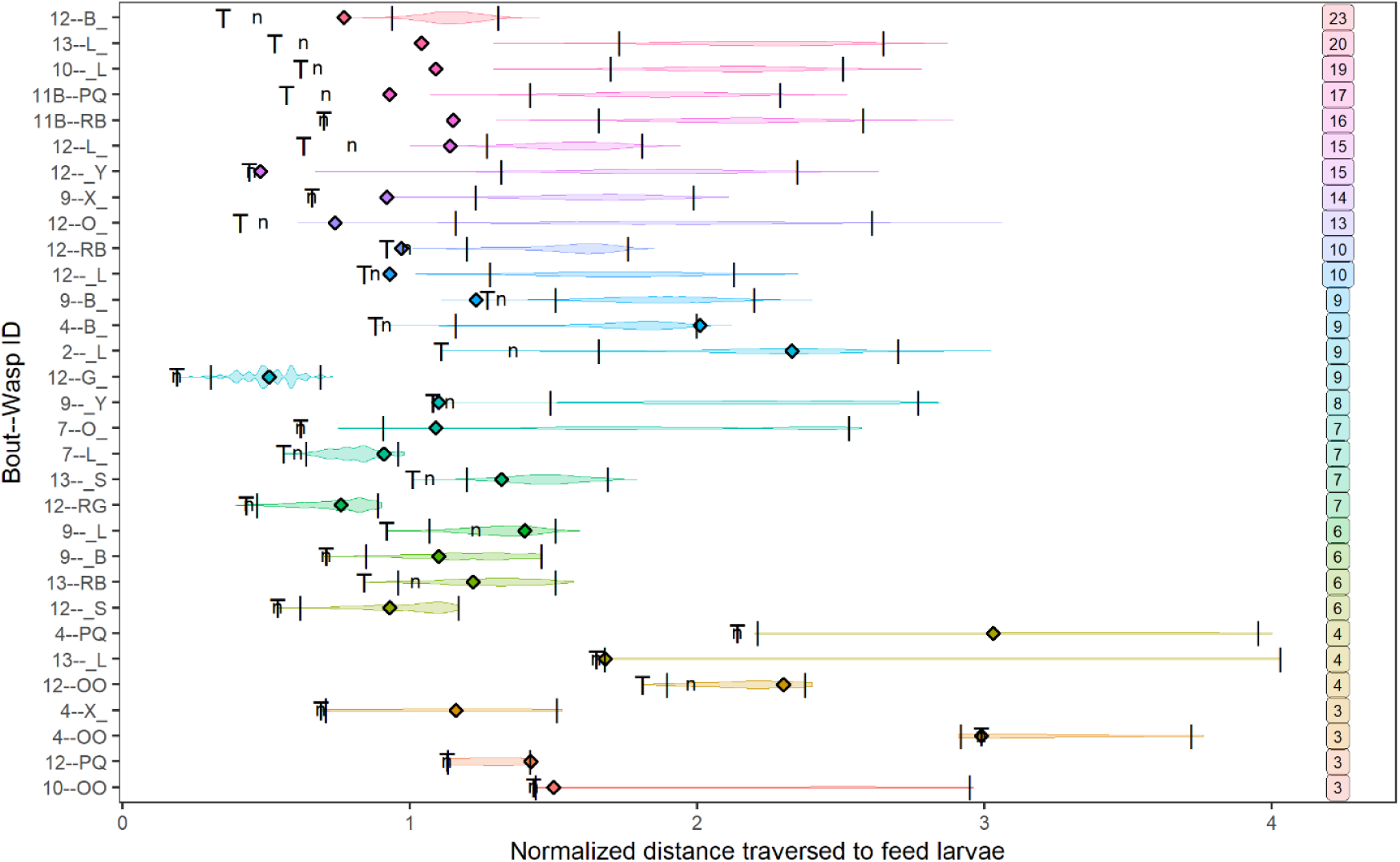
Violin plots showing the distribution of distance per unit larvae fed by an individual within feeding bouts in colony v87. Vertical black lines over each distribution denote the 95% confidence interval. The diamonds denote the observed normalized distance traversed by a wasp while distributing food to the randomly spaced larvae. ‘T’ denotes the minimum possible normalized distance the wasp could have traversed under the travelling salesman algorithm and ‘n’ is the normalized distance it could have traversed by visiting the next nearest larval cell (greedy algorithm). The numbers within boxes on the right are the total larvae fed by each wasp.

**Fig. 6:**
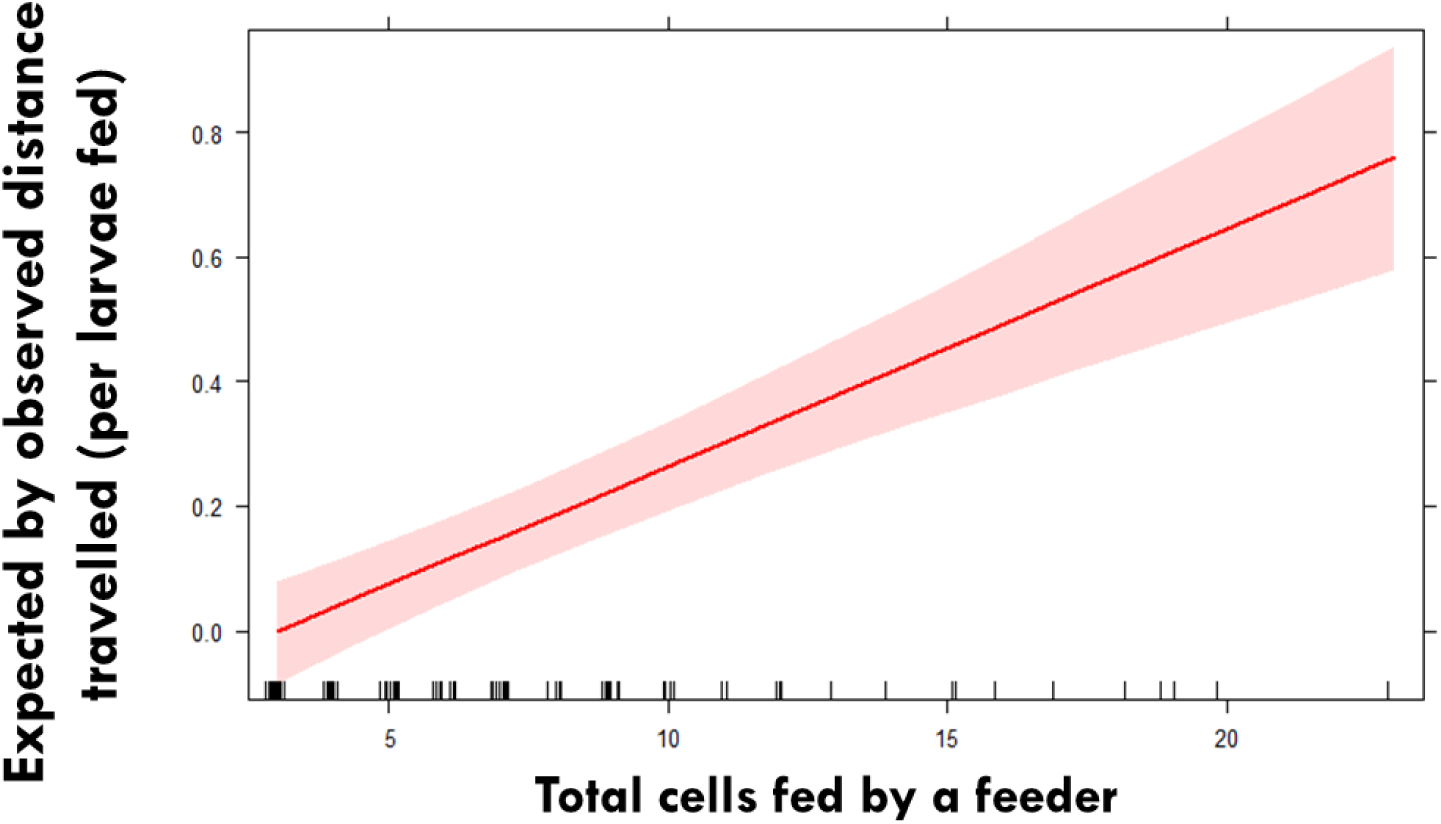
The extent of route optimization measured as the expected by observed distance travelled by a feeder per larvae fed, increased as the number of larvae fed by a wasp increased. Linear mixed effects model was used with log transformed response and wasp ID as random effect.

**Table 2:** The extent of route optimization measured as the expected by observed distance travelled by a feeder per larvae fed, increased as the number of larvae fed by a wasp increased. Linear mixed effects model was used with log transformed response and wasp ID as random effect.

| | Estimate | Std. error | t value | CI (2.5%) | CI (97.5%) | $\chi^2$ | p |
| --- | --- | --- | --- | --- | --- | --- | --- |
| Intercept | -0.11 | 0.05 | -2.12 |  |  |  |  |
| Cells fed | 0.04 | 0.005 | 6.85 | 0.03 | 0.05 | 37.71 | **<0.001**<br>*** |

## Discussion

We studied task partitioning and spatial organization of feeding behavior by wasps within their nests. By spatio-temporally tracking each wasp distributing food within a feeding bout we found that when multiple wasps set out to distribute food in parallel within the same bout, they do not spatially avoid each other on the nest but possibly owing to wasps’ core areas (Sharma and Gadagkar, 2019), when all bouts were put together, the wasps appeared to have larval feeding areas that overlapped less than expected by chance. More interestingly, about a third of the wasps traverse a significantly shorter path than expected by chance, especially when feeding a greater number of larvae within a feeding bout. We also find that wasps preferentially feed larval cells that are closer to the center rather than the periphery of the nest. Additionally, there is considerable inequality in the frequency of larval feeding by primary receivers that unloaded food from foragers.

Division of labour is believed to be one of the leading reasons for the ecological success of social insects. Owing to their morphological indistinguishability, primitively eusocial insects in particular, are known to be totipotent in performing various tasks within the nest (Beshers and Fewell 2001). In our study, in spite of occasional overlap between the performance of feeding larvae behaviour and foraging by the same individual, the two tasks were divided significantly well between colony members.

Across feeding bouts, a majority of nestmates were observed to non-preferentially unload incoming foragers but some receivers unloaded foragers more frequently than others. We delineated Lorenz curves and computed Gini coefficients for each nest to measure inequality in the performance of the unloading task. Gini coefficients are a measure often used in economics to calculate the wealth inequality among citizens of nations or inequality in performance of tasks (Cowell 2000; Crall et al. 2018). Unlike the colonies of *Lasius niger* (Planckaert et al. 2019), foragers in *R. marginata* only feed larvae alone rarely and are often not active participants in food dissemination. In honeybees, the transfer of sub-portions of food load to multiple receivers instead of transferring a larger load to a single receiver allows foragers to get informed about the forager to worker ratio and possibly track the colony demand (Hart and Ratnieks 2001). In *Ropalidia marginata*, we found that about a quarter of the colony members (primary receivers) received food from an incoming forager within a feeding bout. However, the Gini coefficient highlights that there was significant inequality in the frequency of unloading among primary receivers such that some wasps do more unloading than others. We suspect that this differential unloading by multiple wasps could possibly signal some additional information about adult and larval hunger to foragers and could be explored in future studies. There were also no preferred forager-receiver pairs in *R. marginata* nests (as also observed in Planckaert et al. 2019) and while some receivers unloaded food more frequently than others they did not prefer particular foragers to unload food from. We also found that while a majority of primary receivers within a feeding bout either directly fed larvae or acted as ‘conduits’ to transfer nutrition to other adults, about 30% of the primary receivers did neither. Whether these individuals subsequently regurgitate this nutrition or selfishly hoard it needs to be researched further. Inactive workers in other social insects are known to conserve their fat reserves to gain reproductive opportunities subsequently (Lindauer 1952; Jandt and Dornhaus 2011; Charbonneau et al. 2017)..

In *R. marginata*, the presence of randomly located larvae (Sharma and Gadagkar 2019), a lack of clear spatial segregation and high overlap among parallel feeders distributing food to randomly located larvae leads to the central larval cells being fed more than those at the periphery. Such differential feeding frequencies can have important consequences for the individuals as well as the colony. It is known from prior studies on *R. marginata* that individuals that are fed more as larvae, grow to become voracious feeders as adults and are more likely to become future reproductives (Gadagkar et al. 1991). In the nests of bumblebees that also lack a single brood pile, a similar preferential feeding of central larval cells has been identified as a proximate mechanism for shaping adult worker size differences and eventually causing division of labour (Couvillon and Dornhaus 2009). In fact, centralization of resources and information has also been observed in ant colonies where radioactive food was observed to be stocked considerably more in workers located in the colony centre possibly to allow greater access to other nestmates (Aurélie Buffin et al., 2009, 2012; Camazine et al., 2001).

Like in most biological systems including humans and other animals, adopting heuristics for faster problem-solving is often accompanied with systematic errors at individual level that are overcome at the group level (Conradt and Roper 2003; Simons 2004; Surowiecki 2005; Ward et al. 2008; Sasaki and Pratt 2011). Our observation of feeders apparently committing the error of skipping about two hungry larval cells within a feeding bout even though they have food to distribute or feeding the already fed larvae again or, may be an example of such an error committed at individual level that has little cost in terms of larval starvation at the colony level. This becomes evident from our group-level observation where we found a lack of spatial segregation among feeders distributing food to larvae in parallel, within a feeding bout. Repeated visits by different wasps to the same larval cells may in fact be a strategy to overcome such errors made by individuals. This strategy along with retention of food as ‘reserves’ in workers’ crops as speculated in our study, would build in redundancy in the food distribution mechanism such that if some wasps do not receive food in some bouts, larvae do not go hungry. It also prevents cannibalism in extended durations of hunger in natural colonies with sporadic food availability. In the ant *Camponotus yamaokai*, workers show increased trophallaxis during starvation indicating that certain workers retain food in their crops during normal times to regurgitate it during food shortages (Sanada et al., 1998). In the ant *Solenopsis invicta*, workers alter the number of feedings instead of the volume of food to satiate larvae in proportion to their size. An error made by one worker in skipping a larva is made up for, by inspections and subsequent feedings by others (Cassill et al. 1998). While in our study, we allowed foragers to bring food for the colony to mimic a normal colony in nature, it would be insightful to understand if starving the colony or providing *ad libitum* food to the colony influences the tendency of some individuals to regurgitate food in future bouts.

In 27 of the 81 total feeding routes studied (33.3%), wasps distributed food to larvae within each feeding bout by traversing a path that was significantly shorter than random although not the shortest possible path. The extent of route optimization improved as the number of larval cells fed by a wasp increased. This is expected because the benefit from route optimization would be greater when feeding more larvae. Bumblebees are similarly known to improve route optimization with experience when visiting an array of flowers, a behavior called traplining (Lihoreau et al. 2010). We suspect that the wasps which receive bigger chunks of food in the nest are able to feed more larvae and in doing so they traverse shorter routes. Wasps display very rapid, directed movement between larval cells when their mandibles have food (personal observation) possibly in order to get rid of it by feeding larvae and to be available in case another forager arrives with more food in the same feeding bout. We observed a second forager arriving within two minutes (on an average) of the first in almost half of all observed bouts while typical feeding bouts lasted about six minutes (see fig. S4).

Foragers are known to exit in spite of not emptying their crop loads when the unloading rate decreases (Greenwald et al. 2018). As the probability of idle workers taking up additional work is rather low, individuals already engaged in feeding larvae within a bout may be expected to feed larvae quickly to be able to unload a waiting forager (Johnson 2002; Evans 2006; Charbonneau and Dornhaus 2015). Although, it is likely that wasps know the location of larvae within their nest due to frequent patrolling, we do not explicitly suggest that wasps are capable of cognitively computing all possible routes between ‘n’ larvae and then choosing shorter routes. It is possible that much like the extracellular slime serve as memory centres in slime moulds and help select the shortest path to food in a maze (Reid et al. 2012), larvae on the nests of *R. marginata* secrete signals that help wasps to avoid revisiting the already fed larvae – thus reducing their path length. The smell-oriented vertex process proposes the approach to a Hamiltonian path limit cycle if multiple agents follow the same protocol of leaving traces like smell at already-visited sites on a graph even with minimal communication between them (Wagner et al. 1998).

Previous studies have found a role of pheromones instead of behavioral or tactile cues as signals for larval hunger status in colonies of honeybees, bumblebees and ants (Cassill and Tschinkel 1995, 1999; Den Boer and Duchateau 2006; He et al. 2016). Further experiments might help to understand if this mechanism allows wasps to traverse significantly shorter routes by avoiding already visited cells and doing much better than random routing. The observation that most wasps did not follow a TSP, Hamiltonian or greedy algorithm or even a shorter than random route could be because we did not provide any *ad libitum* food in our study and it is possible that there was limited food brought in and most feeders fed fewer larvae in a majority of feeding bouts. As speed-accuracy trade-offs are widespread in the animal world (Franks et al. 2003), we speculate that individual wasps adopt a heuristic to find feeding routes that are shorter than expected by chance especially when carrying enough food to distribute to multiple larvae in the same feeding trip but do not arrive at the shortest path as it might be too cognitively complex or may lead to larger errors leaving larvae hungry.

Several research areas including theoretical computation, mathematics and robotics, derive inspiration from the distributed problem-solving capacity of social insects (Dorigo et al., 2000; Bonabeau et al., 2000; Bonabeau et al., 1997; Krieger *et al*., 2000). Although social insects lack a complex brain, evolution has allowed them to develop alternative strategies to solve mammoth problems within their evolutionary limitations, allowing them to be few of the most ecologically successful animals. Several studies have successfully demonstrated the remarkable computational (Hirsh and Gordon 2001) numerical (Bortot et al. 2019), traplining (Ohashi and Thomson 2009) and transportation capabilities (McCreery and Breed 2014) of ants and honeybees. While a majority of these studies emphasize problems solved in the context of foraging, we propose the new paradigm of studying within-nest food distribution by recording spatio-temporal and behavioural interactions of adults and larvae in detail, to further research on distributed problem-solving in social insects. The wasp system is easily tractable and flexible to manipulations and can have potential applications in the fields of distributed artificial intelligence, communications, navigation and robot mediated disaster management. Biologically, understanding the spatial organization of food transfer may be a key to understanding how insect societies achieve efficient social organization and division of labour.

## Materials and methods

We located three nests of *R. marginata* in and around Bangalore, India (13°00’ N and 77° 32’ E) and transplanted them to the vespiary (A room measuring 10 m×6m×5 m with wire mesh with openings 0.75 cm×0.75 cm instead of sealed windows that allows R. marginata to fly in and out but prevents the slightly larger predatory hornets *Vespa tropica* and *Vespa affinis* from entering) where the wasps were free to fly in and out for foraging. We uniquely marked all wasps with spots of colored paint (Testors® enamel paints - Testors Corporation, Rockford, Illinois 61104, USA) for individual identification. We used video cameras to record all activities on the nest from 8:00 A.M. to 6:00 P.M. (10 hours) for three consecutive days. We collected spatio-temporal and behavioural data for 36 complete feeding bouts. We consider all events of food exchange among adults and between adults and larvae, starting with the arrival of a forager on the nest upto the exhaustion of the food she has brought, as constituting a feeding bout. If a second forager arrives with food before the food brought by the first forager is exhausted, we extend the duration of the bout until all the food is exhausted (see Fig. S1 for representation). We recorded the location and time of each exchange of food among adults as well as between adults and larvae.

### Task partitioning

To measure the extent of partitioning the tasks of bringing food and feeding larvae between wasps, we calculated Gorelick’s division of labour (DOL) index (Gorelick and Bertram 2007) with corrections (Dornhaus et al. 2009), for each nest. To check if the observed values of the index were significantly different from those expected by chance, we compared them with Gorelick’s indices computed assuming all wasps on the nest performed the two tasks randomly, in 100 simulations. To measure the extent of inequality in performing the task of unloading food from the foragers, we calculated the Gini coefficients (using the package ‘ineq’ in R; Zeileis and Kleiber 2014). To check if the observed coefficients were significantly different from those expected by chance, we compared them to coefficients computed assuming random unloading among primary receivers as well as by all colony members, in 100 simulations. Note that we use unloading and receiving food interchangeably throughout the paper.

### Differential feeding of larvae

The sequence of cells inspected and/or fed by each feeder in a feeding bout was recorded. A cell was recorded as skipped if a wasp inspected it but did not feed the larva. Skipped cells could either have been previously fed by another wasp in the same feeding bout or could be skipped even though they had not been fed by other wasps in that bout. Additionally, the proportion of colony members displaying such skipping behavior one or more times was also recorded. The total number of larvae fed by each feeder belonging to three different larval developmental stages (see Fig. S2) was calculated and normalized for the total available cells containing each of the larval stages on the nest. The location of each larval cell on each nest was used to calculate its distance from the colony centre. A linear mixed effects model was used with Poisson error family and larval stage nested in nest ID as random effect to understand if the number of times a larval cell was fed was related to its spatial location on the nest.

### Spatial organization of larval feeding - all bouts pooled

Using the locations (x-y coordinates) of larvae on the nest which were fed in all bouts put together, we delineated the feeding core areas of each wasp defined as the area in which the wasp concentrated 50% of its feeding efforts and computed as the 50% utilization distribution kernels using adehabitatHR (Calenge, 2011) package in R. To test if the core feeding areas of each wasp over all bouts overlapped with each other, we compared the observed average overlap between pairs of feeding core areas of wasps with 1000 simulations assuming random feeding of equal number of larvae by each of the wasps.

### Spatial organization of larval feeding - within bouts

To understand if wasps simultaneously feeding randomly located larvae within the same feeding bout segregate spatially to spread their feeding effort, we simulated 1000 times, larval cells fed by as many wasps as were observed distributing food simultaneously in a bout. Of the total larval cells on a nest, the proportion of cells that were visited uniquely by any one feeder and those visited by multiple feeders were calculated and compared with simulations assuming randomly feeding individuals (see Fig. S3 for cartoon representation).

A feeder setting out to distribute food to randomly located larvae within the nest is analogous to the Travelling Salesman setting out to visit ‘n’ cities. Unlike TSP however, the feeder need not return to the point of origin and this is thus similar to a non-cyclic Hamiltonian path problem. The feeding paths of wasps (see Fig. 6 for example) distributing food to randomly located larval cells on their nests were recorded to understand if wasps solved the difficult Hamiltonian path routing problem by adopting an optimal route or one close to it. As this is an NP-hard problem, whose computation efficiency decreases exponentially as the number of locations visited increases, we exhaustively simulated random routes only for wasps that visited less than 10 larvae (10! = 36,28,800) in a bout. For wasps feeding 10 or more larval cells, we sampled 8000 of the possible sequences of larvae the wasp could have visited. Permutations of larval feeding sequence were calculated using the package combinat (Chasalow and Carey, 2015) in R. The consecutive distance between larval cells for each route was summed to calculate the total route distance. This was divided by the total number of larvae fed by the wasp to normalize and compare across wasps and bouts. For each feeder in a bout, the extent of route optimization was measured as the average expected by observed distance travelled per unit larvae fed by a feeder. We also calculated the shortest possible route the wasp could have taken connecting the larvae it fed as well as if the wasp followed a greedy algorithm (fed the next closest larva based on its current location) by implementing heuristics instead of exhaustively calculating all routes by using the TSP package in R (Hahsler and Hornik 2006). The point of origin in all simulations was retained to be the location of forager alighting with food on the nest. To understand if the extent of route optimization varied based on the number of larvae fed by a wasp, we used linear mixed effects model with log transformed response and wasp ID as random effect.

All the simulations and models were run in R version 3.4.4 (Team R core 2013).

## Supporting information

Electronic Supplementary Material

## Acknowledgements

We thank Priyanka Ambavane for help with data extraction from videos and Kavita Isvaran, Vishwesha Guttal, late Priya Iyer, Ankur Shringi for valuable inputs on analysis.

