## Supplementary material for "Spatial organization of collective food distribution in a paper wasp society": Electronic Supplementary Material

**Electronic supplementary material for:**

| Larval stages | Total FL | Total larval stage on nest | Normalized FL |
|---|---|---|---|
| **L1** | 117 | 47 | 2.49 |
| **L2** | 196 | 44 | 4.45 |
| **L3** | 426 | 72 | 5.92 |

**Table S1**: Number of times larval cells with three different stages of development were fed normalized by the total cells containing the larval stage.

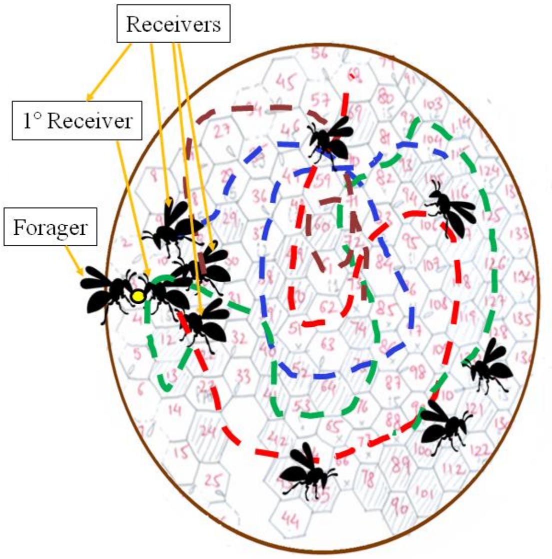


**Fig. S1**: A typical feeding bout involves the forager arriving on the nest with a bolus of food and receivers unloading the food to be distributed further to other nestmates and fed to larvae.

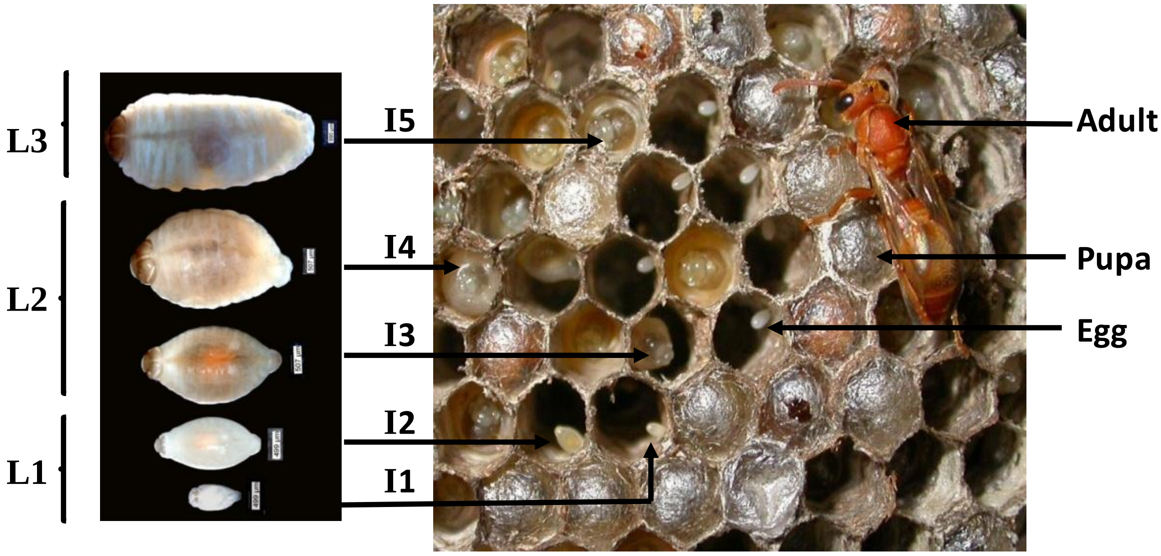


© Thresi and Ruchira

**Fig. S2**: The different stages of larval development in nests of the paper wasp *Ropalidia marginata*

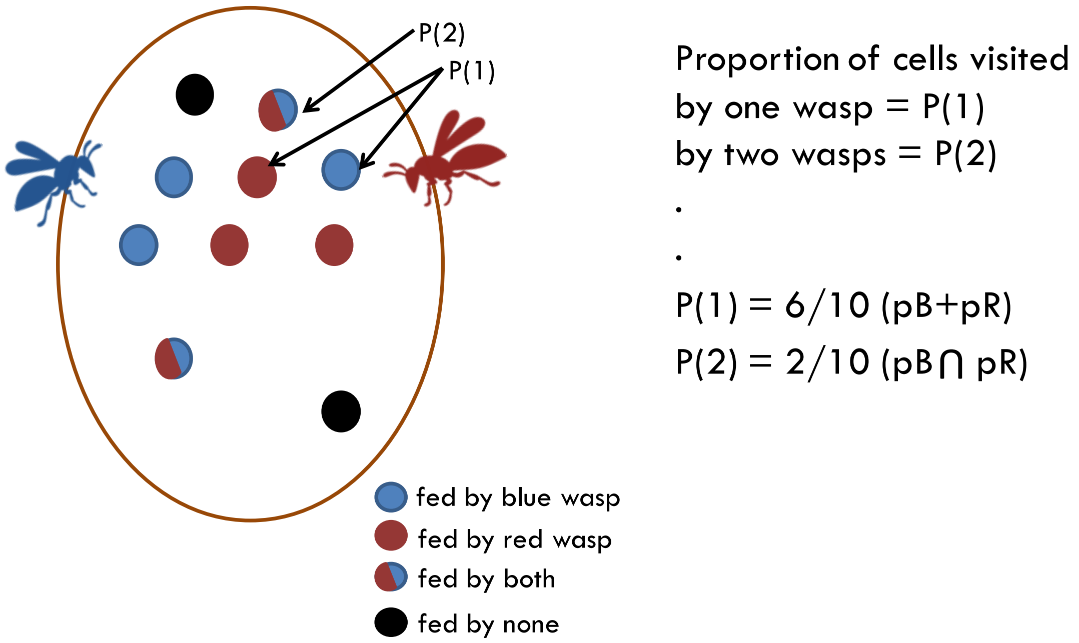


**Fig. S3**: Cartoon representation of the method used to calculate if wasps spatially segregate on the nest within feeding bouts.

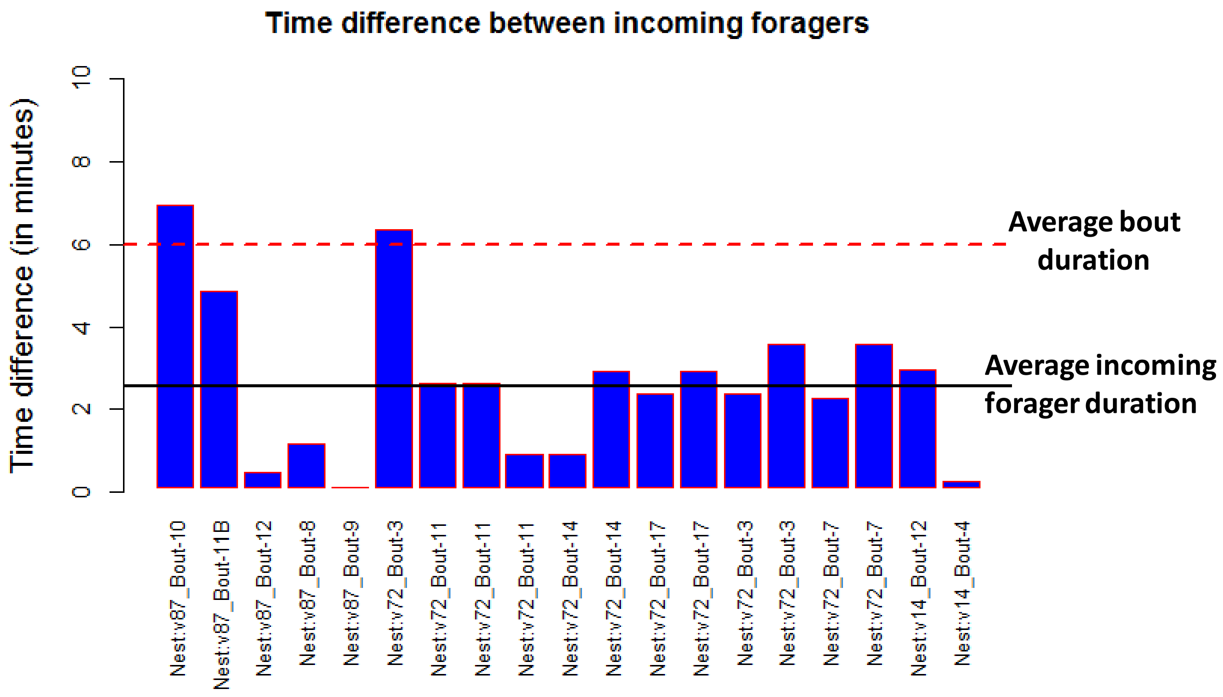


**Fig. S4**: The average time difference between successively arriving foragers in some feeding bouts was 2.25 minutes.

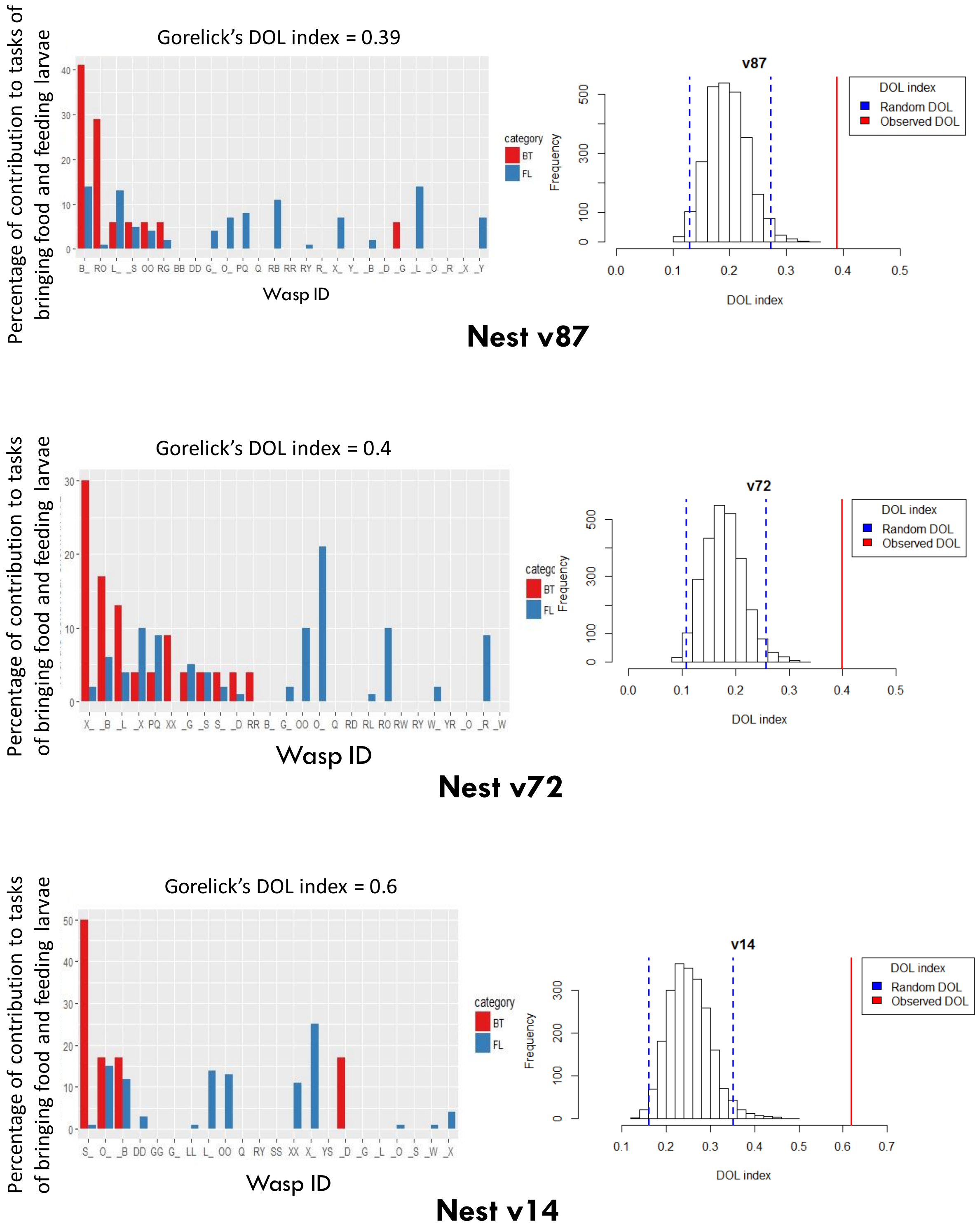


**Fig. S5**: Gorelick's division of labour between the tasks of foraging and feeding larvae was 0.5 on an average and this was significantly greater than if wasps randomly performed either task.

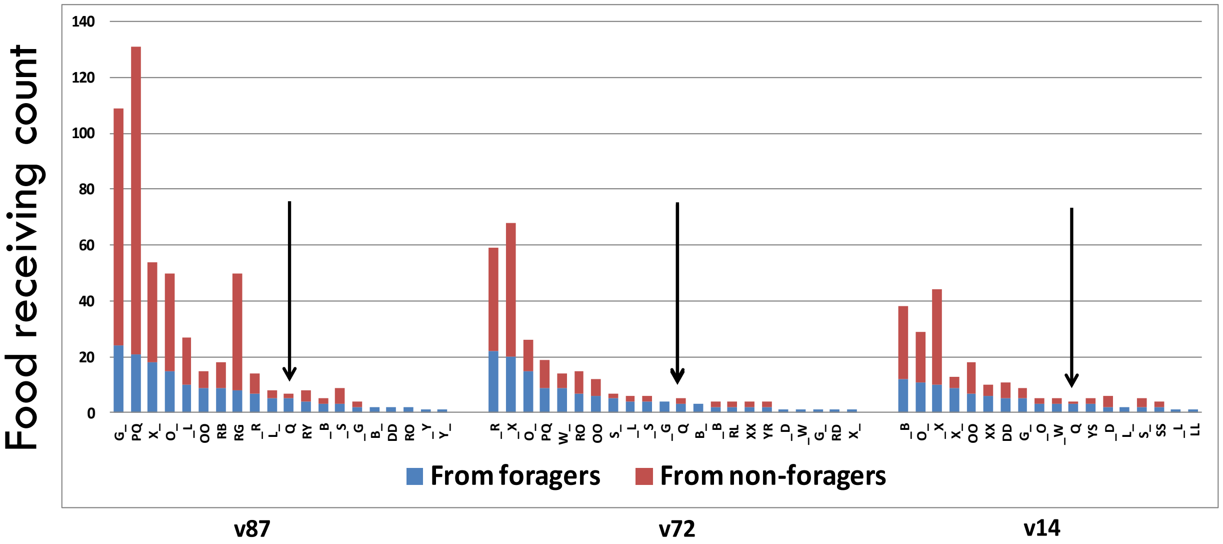


Fig. S6: The queen receives food at intermediate levels

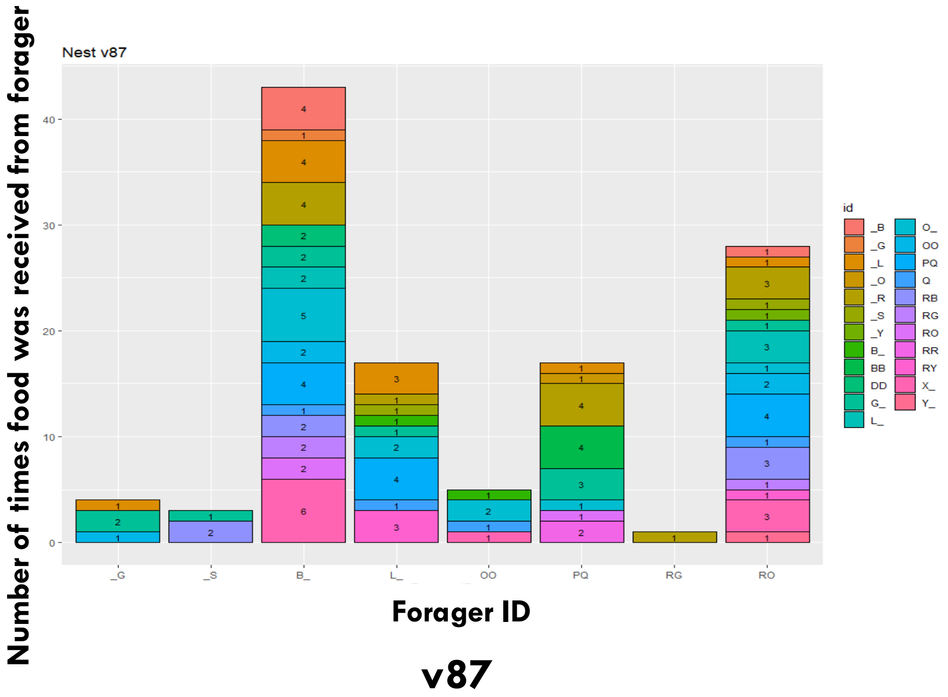

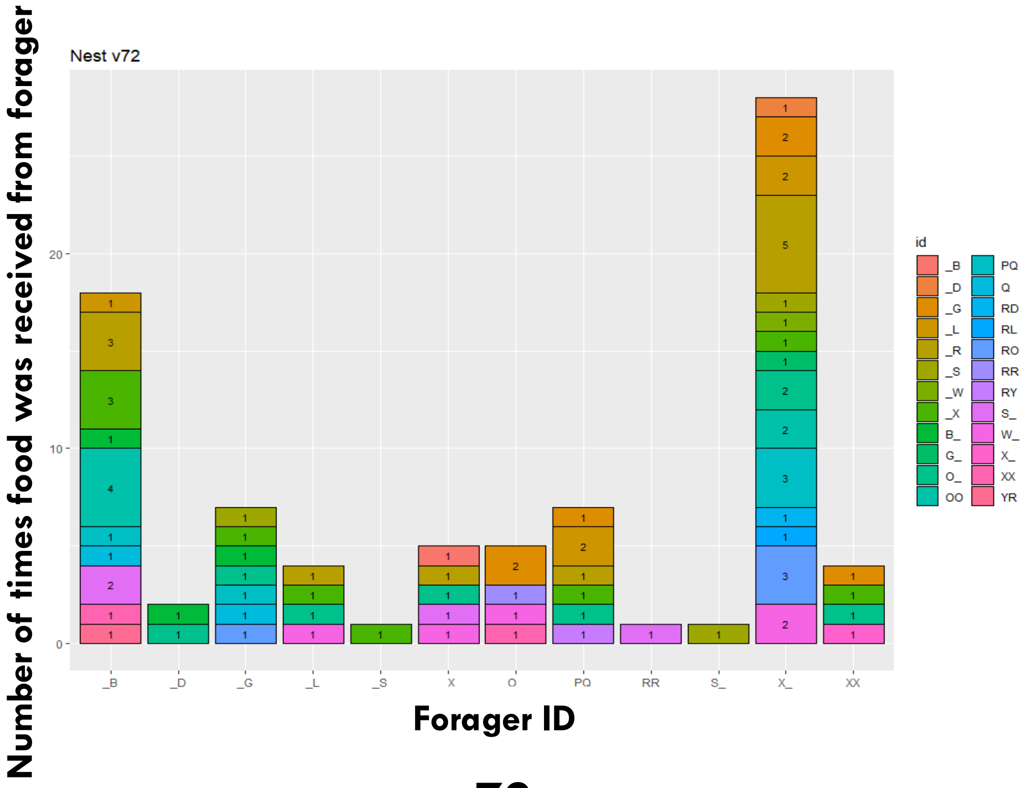


**v72**

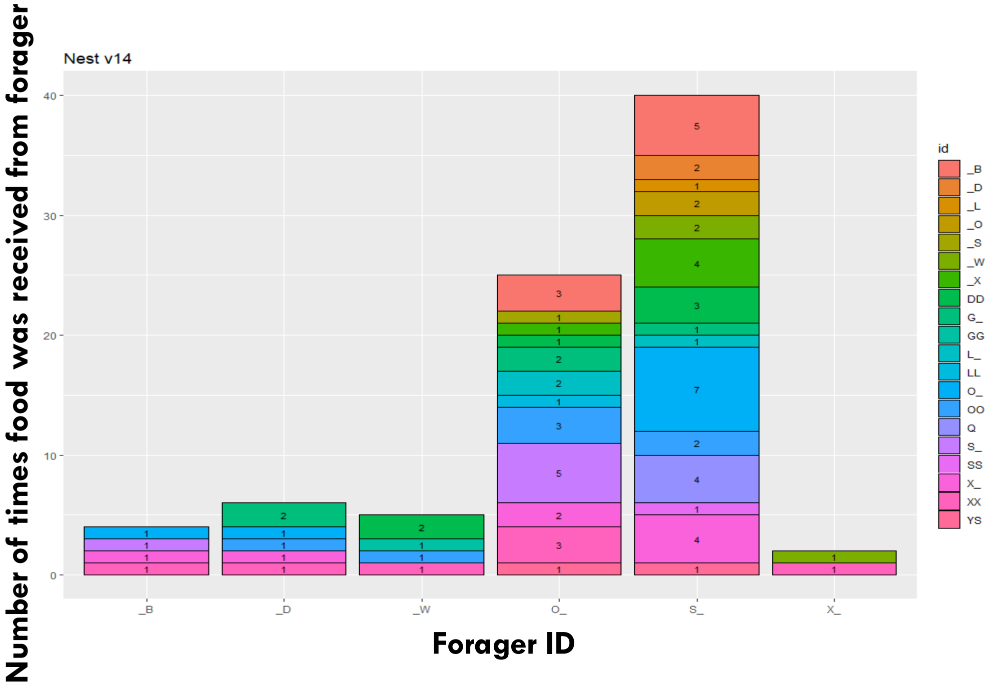


**v14**

Fig. S7: Forager-receiver duos were unlikely

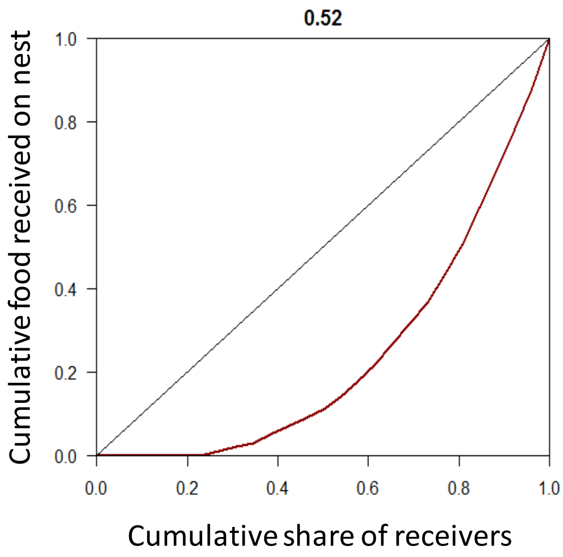
0.52
Cumulative food received on nest
Cumulative share of receivers

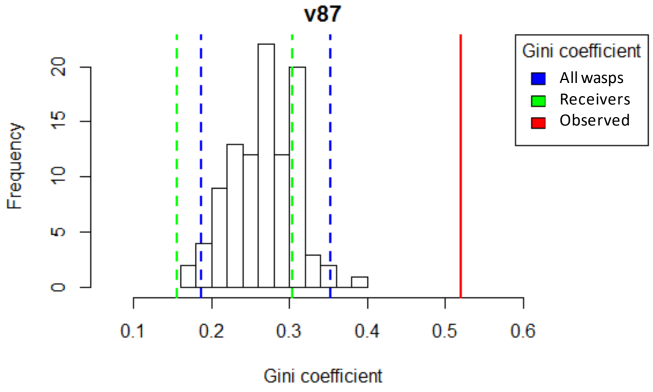
v87
Frequency
Gini coefficient
Gini coefficient
All wasps
Receivers
Observed

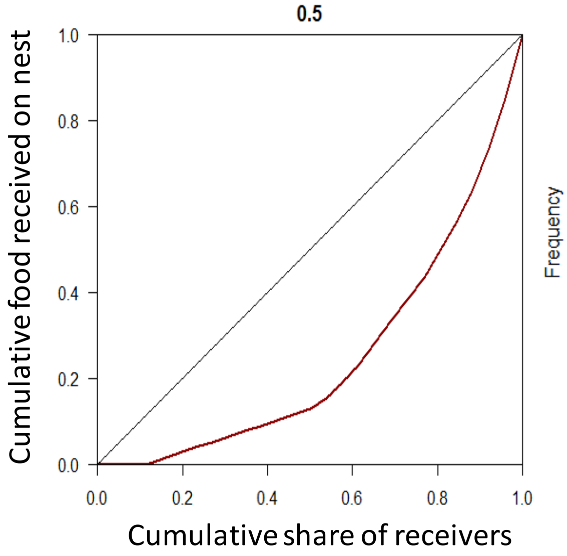
0.5
Cumulative food received on nest
Cumulative share of receivers

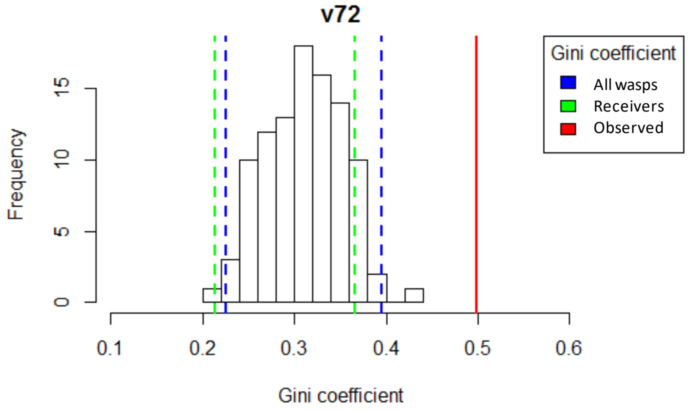
v72
Frequency
Gini coefficient
Gini coefficient
All wasps
Receivers
Observed

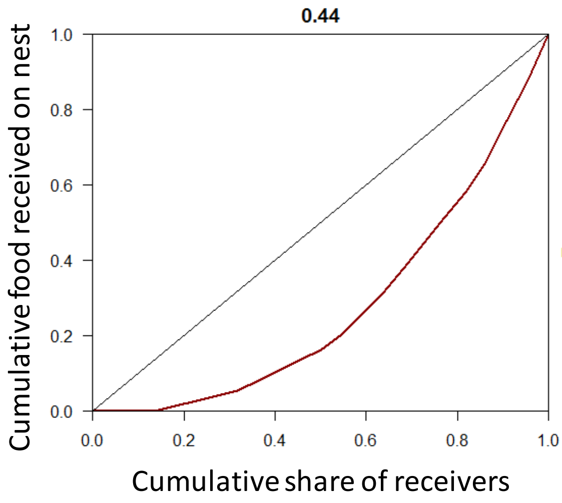
0.44
Cumulative food received on nest
Cumulative share of receivers

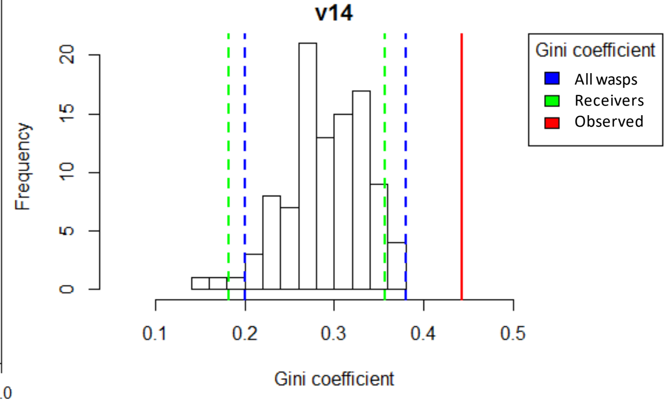
v14
Frequency
Gini coefficient
Gini coefficient
All wasps
Receivers
Observed

Fig. S8: Wasps contributed unequally to the performance of the task of unloading an incoming forager

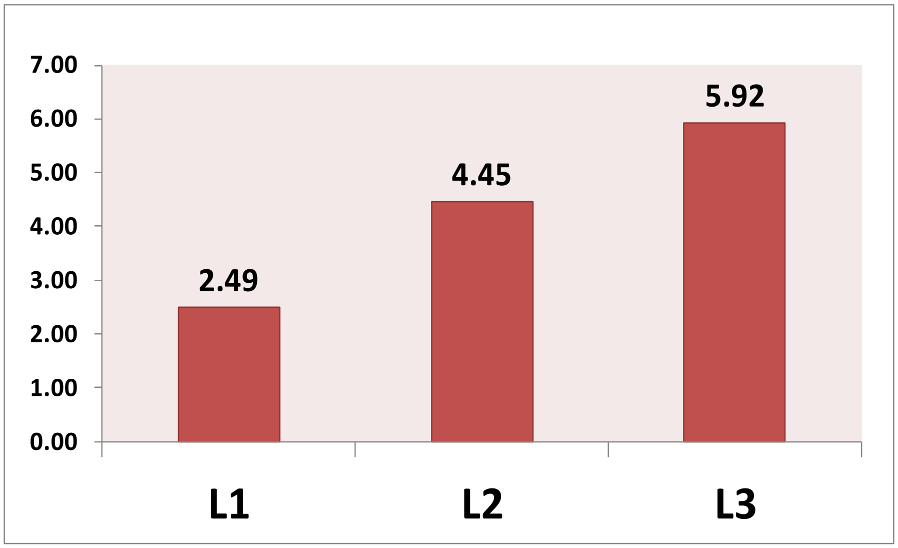


Fig. S9 Pattern indicating larger larvae were fed more frequently

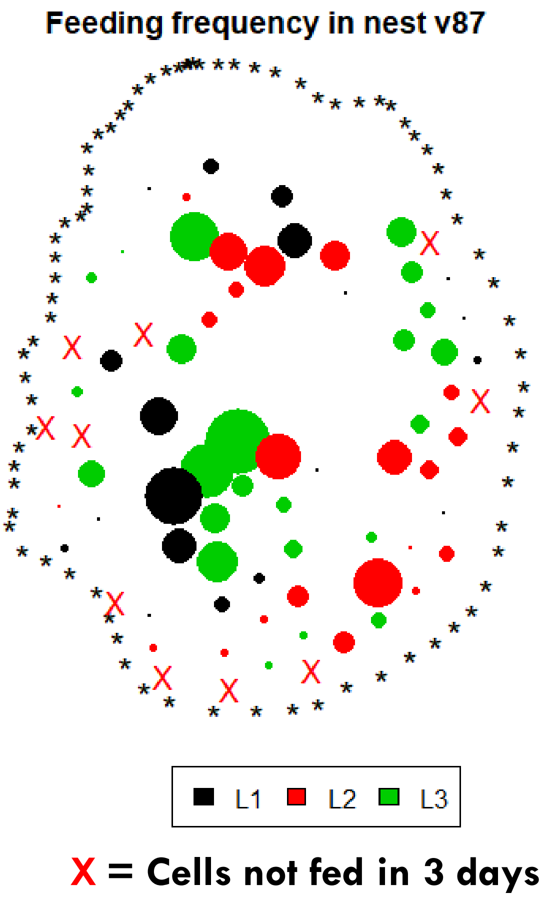


| Larval stages missed | Total cells missed |
|---|---|
| L1 | 5 |
| L2 | 4 |
| L3 | 1 |

**Fig. S10**: Location of larval cells that were fed by wasps in three days of observation of nest v87. The size of the circles is proportional to the number of times that cell was fed. Black boundary indicates the nest boundary.

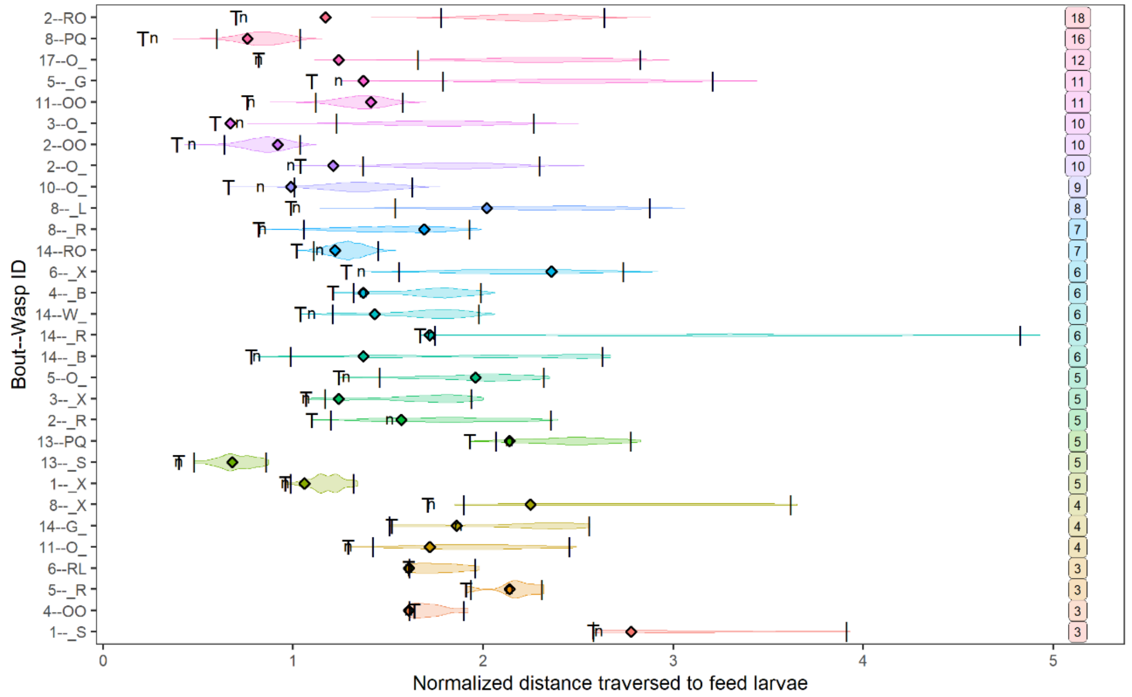


A)

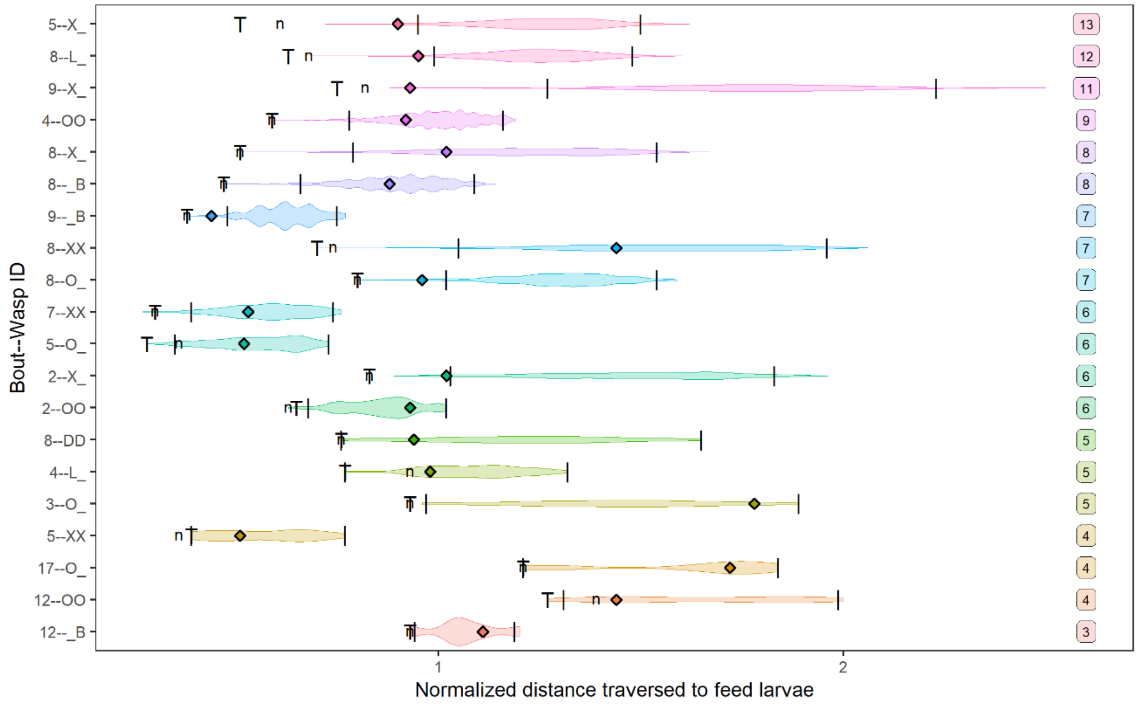


B)

**Fig. S11**: Violin plots showing the distribution of distance per unit larvae fed by an individual within feeding bouts in A) colony v72 and B) colony v14. Vertical black lines over each distribution denote the 95% confidence interval. The diamonds denote the observed normalized distance traversed by a wasp while distributing food to the randomly spaced larvae. ‘T’ denotes the minimum possible normalized distance the wasp could have traversed under the travelling salesman algorithm and ‘n’ is the normalized distance it could have traversed by visiting the next nearest larval cell (greedy algorithm). The numbers within boxes on the right are the total larvae fed by each wasp.
